# Generators of the frequency-following response in the subthalamic nucleus: implications for non-invasive deep brain stimulation

**DOI:** 10.1101/2024.04.30.589054

**Authors:** Mansoureh Fahimi Hnazaee, Haifeng Zhao, Shenglin Hao, Aline Moorkens, Christian Lambert, Shikun Zhan, Dianyou Li, Bomin Sun, Vladimir Litvak, Chunyan Cao

## Abstract

While Deep Brain Stimulation (DBS) is effective treatment for several movement disorders, non-invasive stimulation modes have major clinical relevance. We report on a novel method holding potential for non-invasive subthalamic nucleus (STN) stimulation. We used an auditory frequency-following response task (FFR), a popular tool for studying the auditory brainstem as the neural response in the cortical and midbrain generator, as it precisely reflects the ongoing dynamics of a speech or non-speech sound. We recorded EEG and DBS electrodes from 5 patients, in 4 from the STN, and one from the anterior thalamus and a number of cortical and subcortical areas located in the hippocampus and frontal regions, during an FFR at a frequency higher than the upper limit of phase-locking in the cortex (333Hz). Our results revealed a neural response local to the STN, but not other structures. This finding is novel. Auditory perception in the basal ganglia is rather unexplored, and the STN generator of the FFR has likely gone unseen due to the limitations of our tools and research focus. The potential clinical implications are far-reaching. Future research should investigate whether auditory stimuli at common electrical stimulation frequencies and waveforms of electrical DBS stimulation can induce clinical improvement.

## Introduction

Deep brain stimulation (DBS) is an invasive treatment effective for a variety of brain disorders^1^. Most frequently targeted for movement disorders like Parkinson’s Disease and Dystonia^2,3^ is the subthalamic nucleus (STN), the main input structure of the basal ganglia. In addition to DBS, non-invasive stimulation methods are an important avenue for clinical research^4,5^. Without the need for surgery, they may be more readily embraced and applied at an earlier phase in the disease’s progression. Recent developments are based on temporal interference of electric fields^6^, transcranial direct current stimulation^7^ or transcranial magnetic stimulation^8^.

We report a finding which may hold promise as a novel method for non-invasive STN stimulation. We recorded from STN-DBS electrodes in 4 patients during an auditory steady-state task. This task, the frequency-following response (FFR), is a popular tool for studying hearing. FFRs are particularly suitable for studying the auditory brainstem as they precisely reflect the ongoing dynamics of a speech or non-speech sound^9^. The inferior colliculus (IC) functions as the main site of auditory integration in the midbrain^10^, with multiple studies identifying it as the brainstem generator of auditory responses^11^, using source localization in MEG and EEG^12–16^, lesion studies^17^, as well as invasive neurophysiology in animal models^18–20^. The hierarchically organised structures on the classic auditory pathway act as a low-pass filter for the FFR, such that structures higher up in the hierarchy can lock to the auditory stimulus at a lower maximum frequency. The upper limit above which there is no phase-locking in the cortex is a matter of debate but considered to be ∼200 Hz^13^. A recent study using pure-tone FFR, found cortical contribution to the neural response is not present at 333Hz^16^.

However, it is unknown whether deep structures lying outside of the classic auditory pathway, like the basal ganglia, exhibit FFR and what their phase-locking upper limit is. This partly lies in the limitation of our tools. MEG and EEG source localization studies lack sufficient spatial resolution to distinguish between closely located subcortical nuclei^16,21^. In animal models, the number of electrodes is limited and typically not focused in the basal ganglia^20^. It can also be attributed to the basal ganglia not being of main interest for auditory neuroscience^22,23^. We used a pure-tone FFR at 333Hz^16^, and also recorded the FFR in one patient with implanted electrodes in the thalamus, hippocampus and other subcortical areas. We report FFRs specific to the STN with potential relevance to non-invasive stimulation of the deep brain.

## Materials and Methods

We recorded five patients with implanted DBS electrodes at the Ruijin Hospital of Shanghai Jiao Tong University (see Table 1) after obtaining informed consent (approved by Ruijin Hospital Ethics Committee). The DBS device was from a local manufacturer (SceneRay). Patients were hospitalized for several days after electrode implantation and before implantation of the neurostimulator. During this time, the DBS electrodes were externalized, and we recorded simultaneous local field potentials (LFP) and EEG (64 electrodes, according to the 10-20 system) using Neuroscan (https://compumedicsneuroscan.com/). Sampling rate was 10 KHz. Patients first performed an audiometry screening^24^ to ensure none of the patients had severe or profound hearing loss.

**Table 1.** Clinical Characteristic of patients with implanted DBS in the STN (P1-4) and one patient with depth electrodes in multiple regions (P5)

|  | Age at surgery | Area recorded from | Sex (M/F) | Type of disorder | Medication during recording |
| --- | --- | --- | --- | --- | --- |
| P1 | 44 | Right STN (3 channel pairs) | F | Parkinson's Disease | Levodopa equivalent dose 300 mg/day |
| P2 | 49 | Left and Right STN (6 channel pairs) | M | Muscle Tonic Disorder | None |
| P3 | 55 | Left and Right STN (6 channel pairs) | F | Cervical Dystonia | None |
| P4 | 38 | Left and Right STN (6 channel pairs) | F | Muscle Tonic Disorder | None |
| P5 | 40 | Left and right anterior thalamus, left and right head of hippocampus, right middle and premiddle cingulate, left and right frontal cingulate (108 channel pairs) | F | Epilepsy | None |

The experimental paradigm was a pure-tone FFR at 333Hz similar to^16^, but the interstimulus interval was reduced to 40ms, with no jitter. The trigger for the EEG system was sent 30ms before stimulus onset to prevent electrical interference with the neural response. We recorded 6000 trials of 200ms in alternating polarity per patient (∼24 minutes).

Data was analysed using MATLAB (the MathWorks Inc.). DBS electrodes were localized with lead-dbs^25^. Data was analysed using SPM^26^ and fieldtrip^27^. The data was filtered using a fifth-order high-pass Butterworth filter at 300Hz and a low-pass at 400Hz. No line noise removal was done as the frequency of interest did not overlap with the line noise. Data was cut and trials with an amplitude higher than 4 standard deviations from the mean were discarded. Alternating polarities were subtracted to maximise the response to the temporal fine structure^16,28^. Channels were visually inspected and bad channels were removed. The DBS electrodes were re-referenced to a bipolar montage.

The power-spectra were calculated using a fast-Fourier transform. Inter-trial coherence was estimated by taking the absolute value of the sum of the angle of the Hilbert transform of the signal band-passed at 333Hz±2Hz. Dispersion of the phase was estimated by taking the angle of the standard deviation for the real and imaginary parts of the Hilbert transform separately. Significant FFR response compared to background noise was tested using F-statistics (α=0.5, df=2,11)^29^. Student’s t-tests were done for ITC comparison of EEG and LFP. Bonferroni correction was done for number of tests.

## Results

All patients showed significant FFR in the STN (12/21 STN electrodes, all p<0.005). The electrodes of the epilepsy patient showed a significant response outside the left anterior thalamus (pp<0.0001) and a weak but significant response in the right hippocampus (p<0.0005). No other electrodes survived correction for multiple comparison. Figure 1 shows a summary overview of the results for the phase and amplitude of all patients.

**Figure 1.**
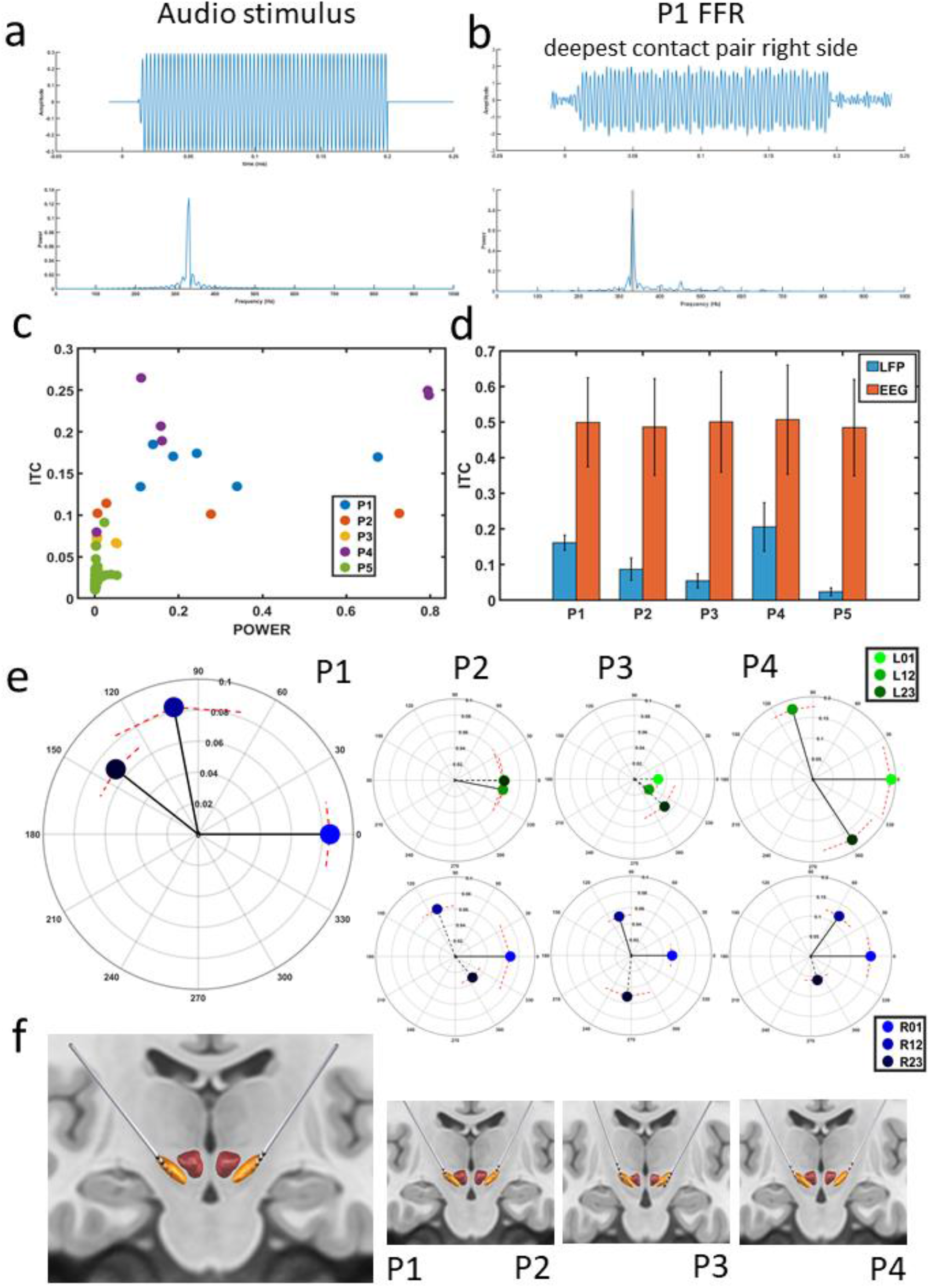
Results of FFR response in the STN of four patients. a) time and frequency domain of auditory stimulus, generated in MATLAB b) time and frequency domain of neural response to FFR for example patient and electrode power, for channel pair optimally placed within STN as confirmed by lead-dbs c) Summary ITC and power of all invasive (LFP) channels, showing power at 333Hz and phase-locking value at 333Hz. d) bar plot for ITC between all invasive (LFP) channels and EEG for each patient. For all patients, the difference was significant at p<0.00001 e) relative phase difference for each STN patient in the left and right hemisphere. The first channel-pair (01) is arbitrarily set at zero phase and phase of other channel-pairs are shown relative to the first. Magnitude of the phase vectors indicates ITC for that channel-pair, dispersion of the phase is shown by red striped lines for each channel pair. Non-overlapping dispersion intervals indicate significant difference in phase difference between pairs. Solid lines indicate a significant FFR response, using F statistics. f) visualization of STN DBS electrodes in the STN, visualized using lead-dbs. Lowest contact pairs are placed within the STN.

## Discussion

We show a response phase-locked to the FFR at 333Hz in the STN. As the STN is anatomically closest to the IC (∼2cm)^30,31^ compared to other channels (closest is the anterior thalamus at ∼4cm^32^), one concern can be that the responses are due to volume conduction rather than generated locally^33–35^. We believe it is the latter based on the following arguments: 1) An obtuse shift in phase, close to a phase-reversal was observed in 5 of the 7 STNs.^36^ 2) Comparing the responses with that of the fifth patient, we conclude they are anatomically specific. The FFR is present in a small proportion of the other electrodes (albeit with smaller amplitude and ITC). Volume conduction cannot explain why some electrodes selectively show the FFR while others in close proximity do not. 3) Finally, the ITC of EEG was significantly higher than the LFP. As LFP is less susceptible to motion and recording artefacts, with pure volume conduction the LFP-ITC would be higher or similar to the EEG-ITC. A reduction in phase-locking could be generated through a functional connection with the IC whereas the EEG is most likely volume conducted from the IC directly. These arguments make a case for a locally generated STN-FFR. Note that we cannot fully rule out volume conduction of a far-field with our methods, one possible way would be to obtain an intraoperative microelectrode recordings of the STN^37^.

Our finding is novel. Speech or auditory perception in the basal ganglia is rather unexplored, with the exception of a few DBS studies recording from the STN and internal globus pallidus^38–43^. At least three of these report on sound perception in the STN, with one using evoked auditory responses showing that wave V, which originates from the IC, appears to pass near the STN^39^. As opposed to neurophysiology, fMRI has adequate spatial resolution to identify all FFR generators^9,44^. While one available FFR-fMRI study analysed only cortical activity^9^, ultra-high field fMRI during auditory stimulation has successfully mapped out brainstem nuclei^45,46^. Mapping brainstem nuclei and connections is challenging due to the densely packed complex structures^47^ that are susceptible to blood pulsation^48^, but whole-brain mappings at 7T compared with conventional 3T^49^ has shown to be promising.

Our finding has implications for anatomical or functional connections between the STN and IC. To date, there is little to suggest direct connections between the STN and IC or other brainstem auditory nuclei. However, there are several lower brainstem motor centres that both directly synapse with the STN and auditory nuclei, including the mesencephalic locomotor region (MLR), caudal pontine reticular nucleus, trigeminal nucleus and nucleus solitaris^48,50–52^. This possible link between the auditory, lower brainstem motor systems and the STN requires further assessment, as it may have possible therapeutic implications. Evidence from possible interactions comes from findings such as “Paradoxical kinesia” in Parkinson’s disease where there is a transient improvement in walking and axial symptoms in response to sudden unexpected visual or auditory stimuli, with the connections between the IC and MLR emerging as a candidate pathway that regulates this motor response^53,54^. Other studies on the auditory startle reflex showed an increase in grip force^55^.

The potential clinical implications of our finding are far-reaching. Future research should investigate whether auditory stimuli at common electrical stimulation frequencies and waveforms of electrical DBS stimulation can induce clinical improvement. Furthermore, this opens new avenues for IC-DBS. Research in animals showed that hearing is not affected by the DBS^56^ but improves tinnitus^57^ and induces motor responses^58^. In light of our findings, studies can extend this to the assessment of clinical symptoms of Parkinsonism.

## Conclusion

We showed a neural response in the STN of 4 patients to the auditory FFR at 333Hz. The response is anatomically specific. Our findings have important implications for future DBS applications by shedding light on the mechanisms of STN and IC functional and anatomical connections. Future studies are needed to map out the potential of auditory stimulation as a novel minimally invasive stimulation method of the basal ganglia, or of IC-DBS for movement disorders.

